# A hormetic model of rapid cold hardening in *Drosophila melanogaster* reveals threshold effects on survival and low fertility resilience

**DOI:** 10.1101/2025.05.12.653569

**Authors:** Jasmine R. Vidrio, Daniel A. Hahn, Michael P. Moore, Gregory J. Ragland

## Abstract

Variable thermal environments may have both detrimental and beneficial effects. For example, extreme temperatures may challenge homeostasis and inflict tissue damage but may also induce acclimation that improves stress resilience. Hormetic models provide a framework to understand dosage-dependent, contrasting beneficial and detrimental effects from physiological and ecological perspectives. We used a hormetic framework and associated quantitative models to investigate how a range of relatively cold, pre-exposure temperatures influence survival and fertility following cold shock at a more extreme cold temperature in the fruit fly *Drosophila melanogaster*. Cold pre-exposure can induce a protective rapid cold hardening (RCH) response, fail to stimulate a response, or cause direct cold injury. We found a plateau-shaped relationship between pre-exposure temperature and female survival resilience, where survival following a cold shock remains high across a range of temperatures, with sharp transitions at higher and lower temperatures. Bayesian fitting of a bi-logistic model highlights these transitions at temperature thresholds that govern processes mediating both acclimation and cold injury. In contrast to survival, female fertility resilience exhibited a muted response to pre-exposure temperature in the presence and absence of post-stress mating opportunities. Overall, a range of pre-exposure temperatures allowed low but successful reproduction following cold shock. High survival but low fertility resilience is consistent with a) differential impacts of cold on somatic and reproductive tissues and b) a growing body of literature suggesting that the thermal sensitivity of fertility may be more limiting than survival for population persistence in variable and changing climates.

**Summary statement:** A hormetic model shows how sharp temperature thresholds govern beneficial rapid cold hardening and detrimental cold injury that have well-defined effects on survival, but only weakly affect fertility.

## Introduction

Variable thermal environments pose a formidable challenge to life, especially when temperatures surpass physiological thresholds for the loss of homeostasis or for induction of cold and heat damage (Schulte, 2014; Williams et al., 2015; Ørsted et al., 2022). On the other hand, variable temperatures also provide opportunities for acclimation, or adjustment of physiology to maintain homeostasis or defend against damage in response to relatively extreme temperatures (Morley et al., 2019; Teets et al., 2023). The net effect of fluctuating temperatures on fitness will depend on the range of temperatures and the duration of exposure, which in turn influence both beneficial processes such as increased cryoprotectant concentrations in preparation for low temperatures (Bale, 1987; Thomashow, 1999) and determinantal or pathological processes such as loss of homeostasis and cellular damage (MacMillan et al., 2015a; Somero, 2020). Many studies explore these beneficial or detrimental effects across ranges of temperatures. However, few investigate how the contrasting effects of these two sets of processes may influence fitness outcomes over temperature ranges that may induce both beneficial and detrimental effects (Yordanov et al., 1986; Williams et al., 2012; Enzor and Place, 2014).

Particularly in ectotherms, nearly all aspects of physiology are plastic as a function of temperature. From an evolutionary perspective, plastic changes can increase, decrease or have no effect on lifetime reproductive output, i.e., fitness (Ghalambor et al., 2007). In general, plasticity that increases or decreases fitness is termed adaptive or mal-adaptive plasticity, respectively. However, most studies only measure one or a few components of fitness, and the same temperature conditions may enhance one fitness component, while negatively impacting others (Kristensen et al., 2008). Thus, whether plasticity is adaptive is often inferred only in the context of one or a few measured traits over some pre-defined, experimental range of environments. Though these complexities mean that strong inferences about adaptive plasticity are difficult to achieve, the general framework of defining physiological impacts of plasticity with respect to fitness is generally useful (e.g., Schulte, 2014)). Here, we avoid the word ‘adaptive’ and consider only whether a plastic response is beneficial or detrimental with respect to specific fitness components.

Whether a given thermal environment increases or decreases fitness may often depend on the combined effects of pathological and protective processes induced by overlapping temperature ranges. For example, a brief exposure to sub-lethal, high temperature may cause protein misfolding, negatively impacting multiple cellular and higher-level physiological processes (Hofmann and Somero, 1995). However, that same exposure may induce the production of chaperone proteins such as heat shock proteins (Hsps) that can mitigate protein misfolding during subsequent exposure to even more extreme high temperatures (Sørensen et al., 2007). Indeed, sub-lethal, short-term exposures to high temperature can simultaneously damage cellular components and also stimulate the classic heat shock response that can be highly beneficial in variable environments (Klepsatel et al., 2016).

Hormesis, a biphasic response to an environmental factor, provides a suitable conceptual and quantitative framework to model these contrasting, beneficial versus detrimental temperature effects. Frequently applied in toxicology, hormesis describes a non-monotonic, biphasic dose-response curve, where a ‘dose’ of some factor (e.g., toxin, radiation, temperature, etc.) increases some measure of performance at low doses, but decreases performance at higher doses (Calabrese and Baldwin, 2001). Beyond toxicology, hormetic models can be valuable in understanding how organisms respond to environmental stressors and can offer insights into ecological processes (Costantini et al., 2010; Berry and López-Martínez, 2020).

Here, we develop and apply a biphasic model to understand the relationship between the intensity of an initial cold exposure and components of fitness measured following subsequent exposure to even more extreme cold in the fruit fly *Drosophila melanogaster* (Fig. 1). Under certain conditions, exposing various insects to moderate duration (often 1 – 4 hours) and moderate intensity (often ∼ 0°C) cold vastly improves survival of a subsequent, more extreme (often ∼ 1hr at β -5°C) cold stress. This short-term acclimation response is known as rapid cold hardening (RCH) and has been well studied as a putative example of adaptive plasticity (or ‘beneficial acclimation’; Woods and Harrison, 2002)) and as a model for rapid physiological adjustments to the environment (Teets et al., 2020). RCH appears to be a widespread phenomenon in arthropods that has also been observed in vertebrates (Teets et al., 2020). Although RCH has often been assessed with respect to survival after extreme cold exposure, RCH also affects other fitness components, e.g., dispersal and reproduction (Lee and Denlinger, 2010; Teets et al., 2020). The initial description of RCH was notable because of the rapidity of acclimation, but also because of the astonishingly strong effects – RCH yielded greater than two-fold increases in survival of extreme cold across multiple insect species, and in some cases increased survival from 0 (no cold pre-treatment) to > 88% with RCH (Lee, et al., 1987). These observations imply a positive relationship between survival and intensity of cold pre-treatment up to a point, conforming to the increasing phase of a hormetic curve where ‘dose’ is the temperature (colder is a higher dose; Fig. 1B). But as the pre-treatment temperature approaches the extreme cold temperature applied in the post-treatment, survival must eventually decline, conforming to the decreasing phase of the hormetic curve. Though the general form of this relationship is self-evident (at least for survival), to our knowledge there are no studies that explicitly model the full function.

**Figure 1.**
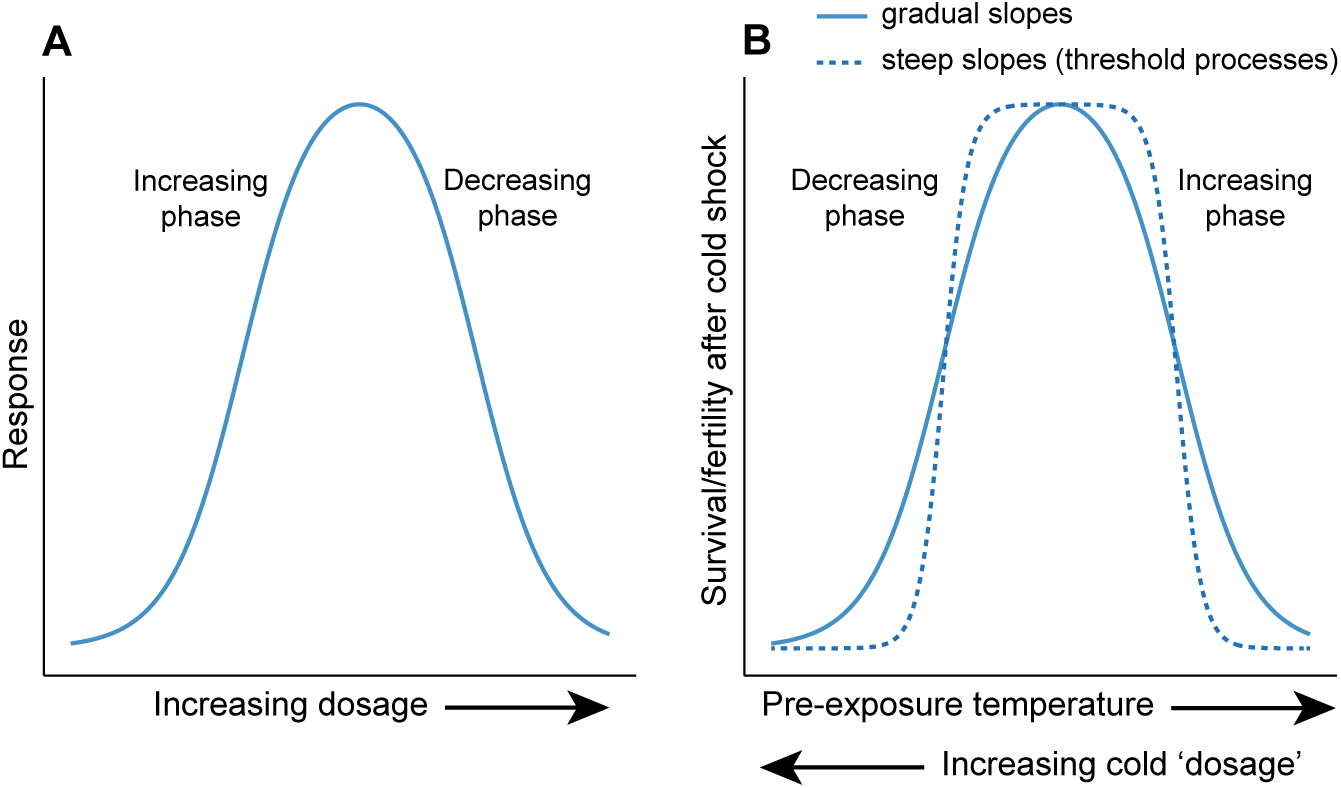
Conceptual diagrams of hormetic models. A) Generic (hypothetical) model depicting a biphasic relationship where some measured response initially increases as a function of increasing dosage (increasing phase) but eventually decreases at higher doses (decreasing phase). B) Hormetic (hypothetical) models relating cold pre-exposure to survival or fertility following a subsequent cold shock. Here, colder pre-exposures represent higher dosages, with initial decreases in temperature increasing survival/fertility after cold shock (increasing phase), and colder pre-exposures eventually decreasing survival/fertility (decreasing phase) after cold shock. The solid line in (B) models gradual slopes, while the dotted line models a relationship wherein processes with tight temperature thresholds underly the observed response.

We performed experiments across a broad range of pre-treatment temperatures to model hormetic curves relating pre-exposure temperature to survival and fertility following extreme cold stress in *D. melanogaster*. This experimental design allowed us to model hormesis in the framework of the standard design for RCH experiments. We also identified an RCH study applying a range cold pre-exposure treatments followed by cold shock to pharate adults of the flesh fly, *Sarcophaga crassipalpis* (Chen et al., 1987). We fit a biphasic model to the results of that study as well to provide a comparison of model suitability and hormetic curve shape across fly species.

Beyond illustrating the utility of biphasic models for understanding both beneficial and detrimental effects of thermal exposures, we had two primary goals.

First, we sought to draw inferences about mechanisms underlying beneficial and detrimental effects of cold exposure. Specifically, we asked whether RCH and cold-damage effects accumulate gradually with temperature, or whether there are tight thresholds for induction. If the physiological effects of temperature accumulate gradually, we would expect shallow slopes relating pre-treatment temperature to fitness components and a relatively narrow range of temperatures over which the fitness component is near the maximum (e.g., Fig. 1B, solid line). If there is a tight threshold for both the induction of the beneficial RCH response and a threshold for when the response to cold transitions from beneficial to detrimental, we would expect steep slopes of increase and/or decrease, and a relatively broad range of temperatures over which the fitness component is near the maximum (e.g., Fig. 1B, dashed line).

Second, we sought to compare the thermal sensitivity of different fitness components, survival and fertility, as a function of pre-treatment temperature. Various aspects of reproduction often demonstrate distinct thermal sensitivity compared to survival and other thermal performance metrics (Walsh et al., 2019). Moreover, survival cannot improve fitness unless successful reproduction follows.

### A note on language and terminology associated with the study of RCH plasticity

We have found that the dual cold exposures, the contingency of measured responses on at least one stressful treatment, and the expectation of a beneficial effect in the name (‘hardening’) can lead to relatively convoluted and sometimes biased language describing RCH experimental results. Below, we simplify the language where possible in three ways. First, we use the term ‘resilience’ to describe the degree of preservation of fitness components (here, survival and fertility) following exposure to a severe stressor (here, severe cold shock). This avoids the need to repeatedly frame the response (e.g., survival) in the context of stressor exposure (e.g., survival following cold shock). Second, we refer to the first temperature exposure treatment simply as a ‘pre-exposure’, which then may or may not improve resilience relative to other pre-exposure conditions. Third, we use the term harden or hardening to specifically describe an outcome in which pre-exposure temperature improves resilience relative to control treatments that do not apply a pre-exposure colder than standard rearing temperature.

## Methods

### Fly lines and husbandry

We performed all experiments on female flies derived from stocks of the isogenic Oregon R *D. melanogaster* line reared in the lab at the University of Colorado, Denver for approximately five years. We used only mated female flies because 1) male and female flies differ in thermal tolerance (Bubliy et al., 2002), and 2) we could easily assay female fertility based on offspring production. We reared all experimental flies in vials on a standard cornmeal–agar–molasses diet in a Percival DR-36VL incubator (Percival Scientific, Perry, IA) set to 24°C and 14:10 light-to-dark cycle.

### Survival experiments

We initially determined that directly exposing flies reared at 24°C to -7°C for 1 hour led to 100% mortality, while pre-exposure to a colder temperature frequently used in other RCH studies (4°C) for 210 minutes led to intermediate mortality following cold shock (30%-35%). Using -7°C as the cold shock temperature therefore allowed us to assess both increases and decreases in mortality with pre-exposures above/below 4°C. All experiments were performed on flies that were 1) anesthetized under CO2 and sorted into new vials with food 3 – 4 days post-eclosion, then 2) allowed 24 hours to recover from anesthesia, then 3) tap-transferred into empty vials. Using the direct transfer RCH method from Gerken et al. (2018), we transferred fly vials from the rearing temperature of 24°C directly into an Arctic A40 recirculating chiller (ThermoFisher Scientific, Waltham, MA, USA) containing 50 % (v/v) propylene glycol that was set at the desired pre-exposure temperature. Vials were held at the pre-exposure temperature for 210 minutes, then the bath temperature was dropped at a constant rate over 5 minutes to the acute cold shock temperature of -7°C, then held there for 60 minutes. To monitor the bath temperature, we placed two Type T thermocouples (Omega Engineering, Norwalk, Connecticut, USA) interfaced with PicoLog v6 via TC-08 units (Pico Technology, Cambridgeshire, UK) into each of two empty vials (four thermocouples total) placed proximal to vials containing experimental flies. Vial temperatures were always within 0.22°C of the intended pre-treatment/cold shock temperature (see data archive, Data availability statement). Following the cold shock, flies were gently tapped into new food-containing vials and immediately transferred back to 24°C. Vials were placed onto their sides during transfers and for 24 hours following transfers to avoid comatose flies sticking to food. After 24 hours, flies were scored as “live” if they displayed coordinated movement and were able to stand on their legs (David et al., 1998).

We applied a broad range of pre-exposure temperatures, attempting to identify a range of temperatures causing high survival resilience, and temperatures above/below that range causing relatively low survival resilience. In initial pilot experiments, we also noted a rapid decline in survival with pre-exposure temperatures below 0°C and thus applied more closely spaced pre-exposure temperatures at the low end of the pre-exposure temperature range. We applied the following pre-exposure treatments: 210 minutes at –1, –0.5, 0, 2, 4, 6, 8, 10, and 14°C. Figure 2 illustrates the thermal profiles for each treatment – treatments varied in pre-exposure temperature but all included a common 60 minute exposure to a -7°C cold shock. Each treatment included three to four replicate vials (see data archive, Data availability statement), with each vial containing 20 females.

**Figure 2.**
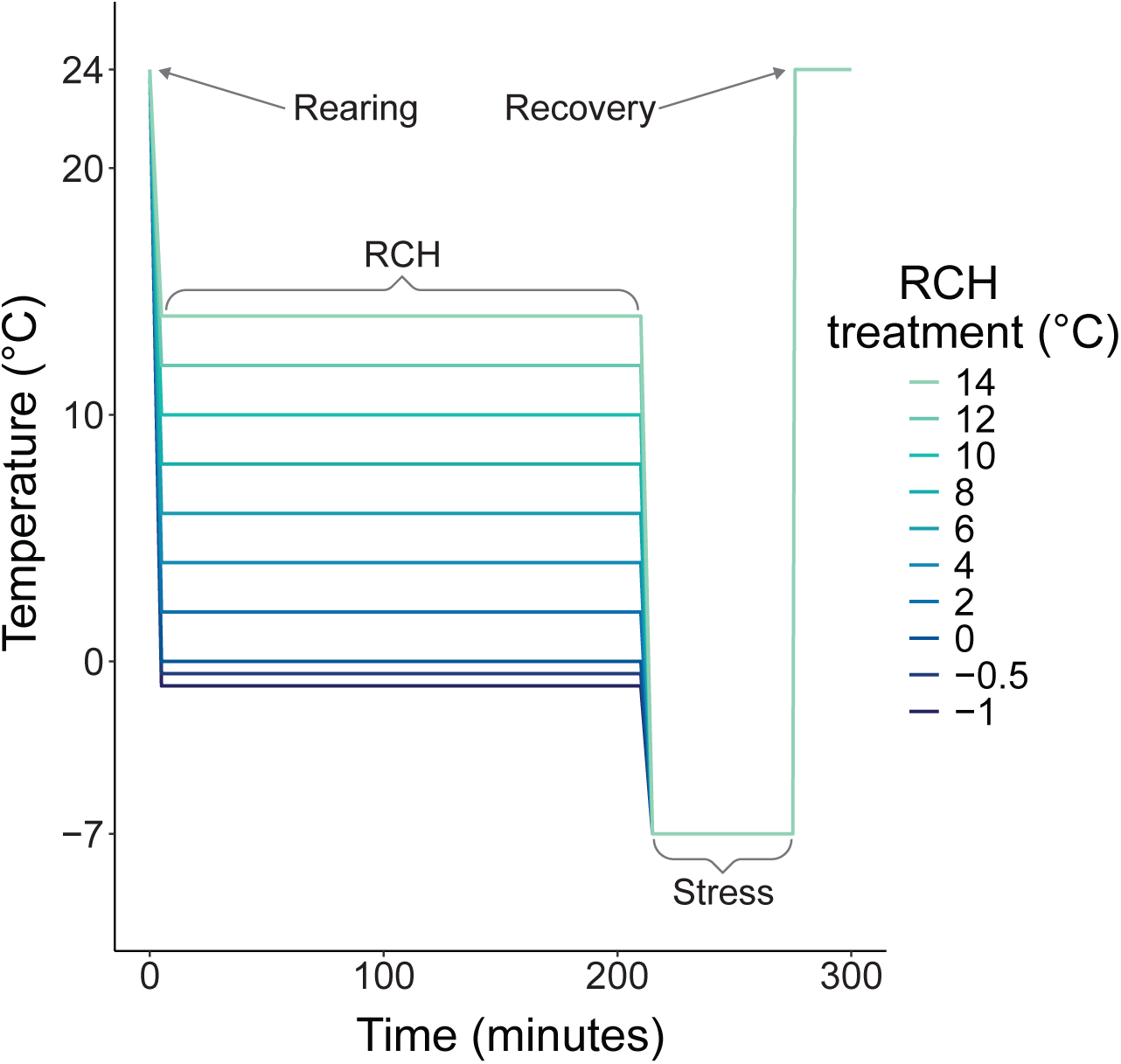
Conceptual illustration of experimental treatments. Each treatment (line) used common rearing, cold shock, and recovery temperatures, with only pre-exposure temperature varying among treatments.

### Fertility assays

Following the survival assays described above, we tracked offspring production of surviving females. Counting eggs is a common method to assess fertility, but eggs may or may not be viable, and offspring may or may not successfully develop. We instead counted the number of pupae produced in each vial as in, e.g., Hafezi et al. (2020). Females (without any males added) were allowed a total of 48 hours following the cold shock exposure to oviposit and then were cleared from the vials. We also created control vials containing 4 -10 three-to-four-day-old females from the same cohort of flies used to initiate the experiments. Control flies were handled the same way as pre-exposure/cold shock treated flies but were simply held at the rearing temperature (24°C) for the duration of treatment period. Recall that we used mated females that can continue to produce viable offspring from eggs fertilized prior to the experiments or using stored sperm. We counted total pupae produced from each vial up to 14 days following the experiment – no new pupae were ever observed past 14 days, much longer than the usual egg-to-pupa development time for *D. melanogaster* at 24°C.

After initially observing very low fertility across all pre-exposure treatments (see results), we performed one additional experiment to test whether introducing untreated males to females after they had been cold shocked might improve fertility resilience. We repeated the assays as described above applying two pre-exposure temperatures, one where survival was relatively high (10°C pre-exposure) and one where survival was relatively low (14°C pre-exposure). We also included untreated control vials, as above. We modified the assay by splitting vials into two groups: one group had no males added (equivalent to the treatments described above), and one group had 20 untreated, non-virgin, 3-4 day post-eclosion males added per vial immediately following the return to 24°C following cold shock. Vials were cleared at 48 hours, and pupae were counted as described above. Each pre-exposure/cold shock treatment included four vials with and four vials without females, respectively, while each control treatment included one vial. If fertility in this experiment recovered after untreated males are added to the vials, the result would suggests that pre-exposure and/or cold shock is disrupting reproduction through its effects on the male contributions stored by females prior to their cold shock, perhaps through reduced sperm motility or viability. By contrast, if fertility did not recover after introducing untreated males to cold shocked females, the result would suggests that pre-exposure and/or cold shock disrupts other aspects of female reproduction through, e.g., damage to somatic or germline reproductive tissues.

### Statistical analyses

Hormetic processes can be modeled using a bi-logistic function including both increasing and decreasing logistic phases (Beckon et al., 2008). Such functions can include easily interpretable parameters that describe, e.g., the rate of increase/decrease, thresholds or positions of phases (often inflection points), and maximum response values (e.g., survival or fertility in this study). Initial inspection of the data suggested a non-monotonic, plateau-shaped function with asymmetric rates of increase and decrease. We designed a modified logistic, 5-parameter model to incorporate these shape elements. Our model is a simplified variant of the biphasic model described in Beckon et al. (2008). This simplified model is is less flexible in shape, but it provides a good fit to our survival data (see results) and has the advantage of more straightforward parameter interpretation. We modeled the relationship as follows:

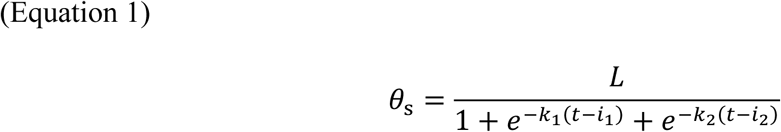

where θ*_s_* is the proportion survival, *t* is the temperature, *L* models the upper limit of survival, *k*_1_ and *k*_2*_model the steepness of logistic change in the decreasing and increasing phases, respectively (Figure 1B.), and *i*_1_ and *i*_2*_ model the temperatures of the two inflection points (the temperature at which the logistic changes in each phase are centered). Note that the codomain of the function is any real number; thus, it can be used to model data in different ranges (e.g., [0,1] for survival proportion) by constraining *L*. We fit the model using a Bayesian procedure implemented in Stan (6.1 Stan User’s Guide Version 2.34) and accessed through the *cmdstanr* package in R version 4.3.1 (R Core Team, 2023). We used the default No U-Turn Sampler (NUTS), a Hamiltonian Monte Carlo (HMC) method to sample the posterior distribution, running 4 sampling chains, each with a burnin of 2000 iterations followed by 3,000 retained samples per chain, 12,000 samples total. Because survival proportion in some treatments was well above 0.5, we constrained *L* to [0.5,1] and we placed additional constraints and priors on some parameters based on visual inspection of the data to prevent non-convergence (see relevant scripts in the Zenodo archive and supporting file 1 for details and rationale). We specified a binomial (*n* = number of total females in a vial, *k* = number of surviving females) likelihood function for θ*_s_*. All chains achieved stationarity with symmetrical, unimodal posterior distributions for each parameter. Finally, to visualize the modeled relationship with uncertainty, we calculated mean θ*_s_* and the 95% credible interval of the posterior distribution for temperatures ranging from -3 to 18°C at 0.01°C increments.

To assess whether the model fit well and predicted a similar relationship for a comparable set of data from another insect species, we fit the same bi-logistic function to estimates of the survival resilience to cold shock across pre-exposure temperatures reported in a separate study of *S. crassipalpis* pharate adults (Chen et al., 1987). We extracted estimated mean proportion survival vs. pre-exposure temperature from Figure 3 in Chen et al. (1987) using PlotDigitizer (https://plotdigitizer.com/app). We set the binomial parameter *n* to 60 for each trial (the study methods indicate 15 – 20 pupae in each of three replicate groups) and we set *k* to produce the estimated proportion survival reported in Chen et al. (1987). We did not estimate uncertainty (the 95% credible interval) for the model using the data from Chen et al. (1987) because we lacked raw data including survival rates per replicate group and exact sample sizes.

**Figure 3.**
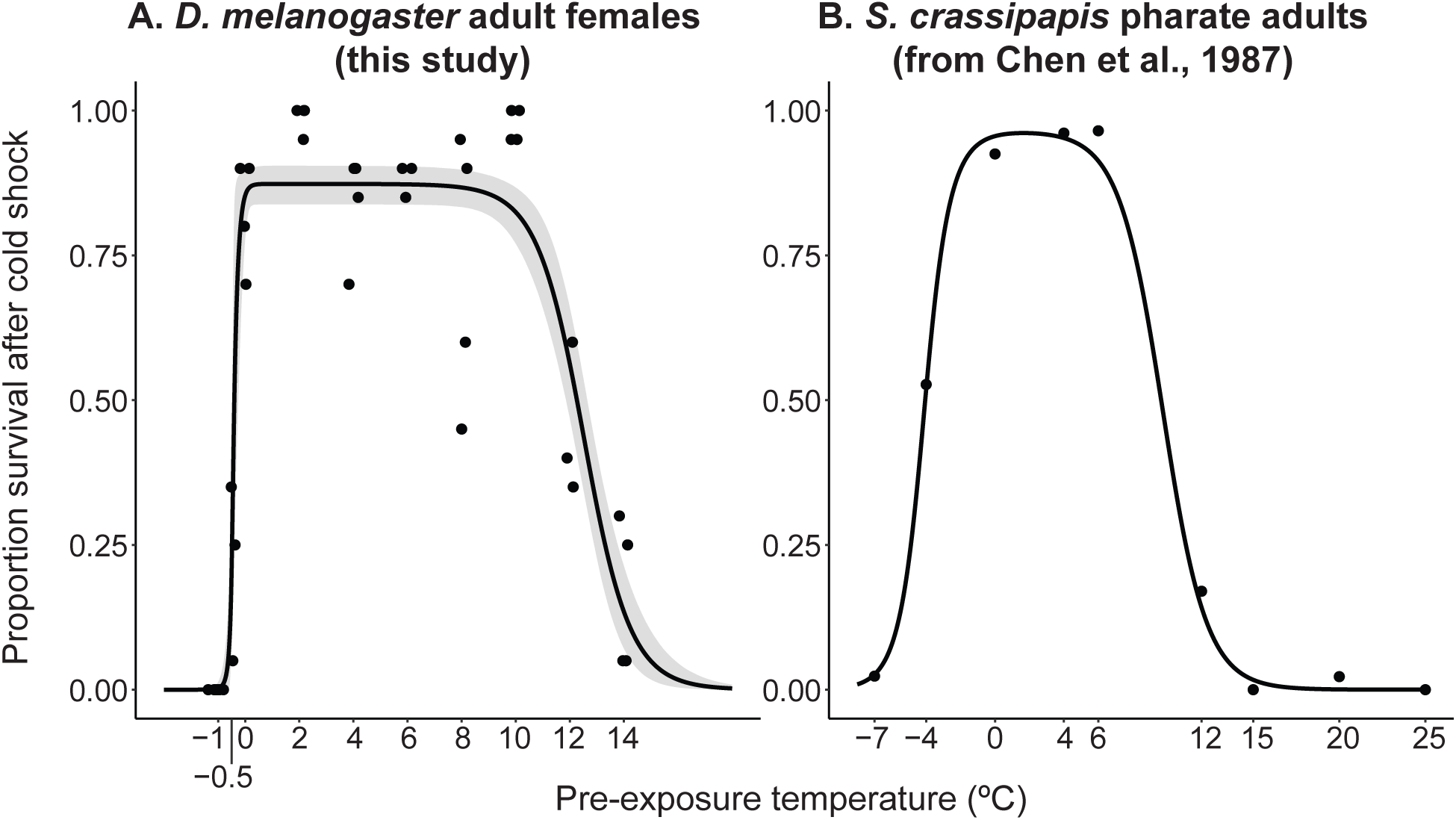
A) Survival resilience (post cold shock) of 4-5 day old female D. melanogaster vs. pre-exposure temperature. All flies were held at the pre-exposure temperature for 210 minutes followed by 60 minutes of stress exposure at -7°C. The black solid line and the grey shaded area are the fitted bi-logistic model and the 95% CI, respectively; filled black circles are the mean proportion survival of each replicate vial, jittered horizontally to limit overlap. B) Survival resilience of pharate adults of S. crassipalplis vs. pre-exposure temperature. Filled black circles are mean proportion survival estimated by Chen et al. (1987); the solid black line is the fitted bi-logistic model (CI not estimated because the model was not fitted to raw data).

Initial inspection of the first set of fertility data, including all pre-exposure temperatures from the survival experiment and without addition of males, suggested low fertility with high variance within temperature treatments. Attempted fitting of a modified version of the model described in equation 1 (using a negative binomial likelihood function) resulted in non-convergence with moderate constraints and priors comparable to the survival model above. We were able to fit the model with highly constrained parameter values and highly informative priors, which we present in a supplementary file. However, the model fit was relatively poor and overly influenced by constraints and priors and we provide the details in the supplement only as a proof of concept that the model can be fit to different data distributions.

We chose to instead fit generalized linear models (glm) treating pre-exposure as a discrete factor to simply test whether fertility varied across pre-exposure temperature using the *glm* function in R. Though glms with Poisson and related error distributions require count data as the response, rates can be modeled by including an offset term (Dunn and Smyth, 2018). We used this approach to model the rate of pupa per female using glms with total pupa per vial as the response, pre-exposure temperature as a discrete factor, and number of surviving females per vial as an offset. An initial fit using a Poisson model suggested overdispersed data using the simulation-based dispersion test from the *DHARMa* package in R. We settled on a negative binomial model with the same parameters and data, providing a good fit with no evidence of overdispersion. We then used the *predict* function to estimate mean pupa per female and associated 95% confidence intervals (CI) for each pre-exposure and control treatment.

The second fertility experiment included only two pre-exposure temperatures and a discrete factor for addition (or non-addition) of males. We thus applied a similar approach using negative binomial models with total pupa per vial as the response and number of surviving females per vial as an offset. We fit two models – both included a discrete factor for pre-exposure temperature and a discrete factor for male addition, but one also included an interaction. We used AIC model selection to determine the best fit model, then used the *predict* function to estimate the mean pupa per female and associated 95% CI for each treatment combination.

Finally, we estimated replacement rate under laboratory conditions to illustrate how cold pre-exposures influence a composite metric of fitness including both survival and fertility after cold shock. For vials that were measured for both survival and fertility (all vials in the survival experiment that were also measured for fertility without the addition of males), we estimated laboratory replacement rate of the females as the product of survival proportion and fertility (pupae per female) estimated for each replicate vial. The variance in replacement rate within and among treatments appeared similar compared to estimated fertility. Thus, we did not attempt to fit a continuous, Bayesian model. Rather we estimated means and approximate 95% confidence intervals (mean ± 2SEM) for replacement rate at each pre-exposure treatment.

## Results

### A broad hormetic curve with asymmetric slopes relates survival resilience to pre-exposure temperature

The bi-logistic model fit the data well and suggested a broad range of temperatures over which pre-exposure to moderately low temperature improved survival resilience to cold shock (Fig. 3A; Table 1). Low survival resilience with a pre-exposure of 14°C suggests that higher temperatures are inadequate to stimulate a hardening response. The slope of the increasing phase at higher pre-exposure temperatures was relatively steep, with the mean predicted survival increasing from 14% to 83% survival over 4°C from 14°C to 10°C pre-exposure treatments. This pattern is consistent with a threshold process wherein decreasing pre-exposure temperature lower than 10°C (and therefore applying a more intense pre-treatment) provides no greater protection from cold shock induced chilling injury. Though there was substantial variability (illustrated by the variation in vial means), predicted mean survival resilience remained similarly high from pre-exposures between 10 – 0°C, quite a broad range of temperatures that induced similar resilience.

**Table 1.** Parameter estimates for the bi-logistic model fit to D. melanogaster survival data including the mean, 0.05 quantile, and 0.95 quantiles of the posterior distributions.

| Parameter | Posterior Mean | 90% CI |
| --- | --- | --- |
| $k_1$ | 16.3 | 6.86 - 37.0 |
| $k_2$ | -1.17 | -1.47 - -0.89 |
| $c_1$ | -0.407 | -0.475 - -0.313 |
| $c_2$ | 12.5 | 12.2 - 12.8 |
| $L$ | 0.873 | 0.844 - 0.900 |

Survival resilience declined precipitously at pre-exposure temperatures below 0°C, dropping from mean predicted survival of 85% to 0% survival from 0°C to -1°C. This suggests a very tight threshold for either chilling damage, inhibition of physiological processes that underlie hardening, or some combination of both. The fitted model predicts survival resilience in this temperature range that is higher than zero (the outcome with no pre-exposure) yet much lower than the survival resilience induced at the plateau of the hormetic curve. Thus, 0°C to -1°C is a pre-exposure temperature range inducing a combination of both limited hardening and chilling damage.

### A similar hormetic curve relates survival resilience to pre-exposure temperature in the flesh fly Sarcophaga crassipalpis

Data from Chen et al. (1987) reveal a very similar relationship between pre-exposure temperature and survival resilience in a different life stage (pharate adults) in a different fly species (*S. crassipalpis*). The fitted bi-logistic model suggested clear pre-exposure temperature thresholds bracketing a range of pre-exposure temperatures over which survival resilience was similarly high (Fig. 3b), albeit over a narrower range than for *D. melanogaster* adult females.

### Pre-treated, surviving flies had universally low, but non-zero fertility

In contrast to survival, fertility resilience was universally low compared to fertility in control flies that did not experience pre-exposure cold or cold shock (Fig. 4A). There was a weak relationship with pre-exposure temperature, with fertility slightly higher from 6 - 10°C. This pattern hints at a hormetic relationship similar to survival, but the outcome was highly stochastic. For example, fertility was zero in the 0°C treatment despite reasonably high survival, but non-zero in the -0.5°C treatment where survival was low. Addition of unshocked males back to groups of cold shocked females to allow for post-recovery mating had only marginal effects on fertility resilience, and only in the 14°C pre-treatment (full model, including interaction AIC = 102.6; reduced model excluding interaction AIC = 126.2) in which fertility resilience was moderately higher with the availability of males compared to no males (Fig. 4B). However, fertility resilience of all flies in pre-exposure plus cold shock treatments was much lower than fertility of control flies.

**Figure 4.**
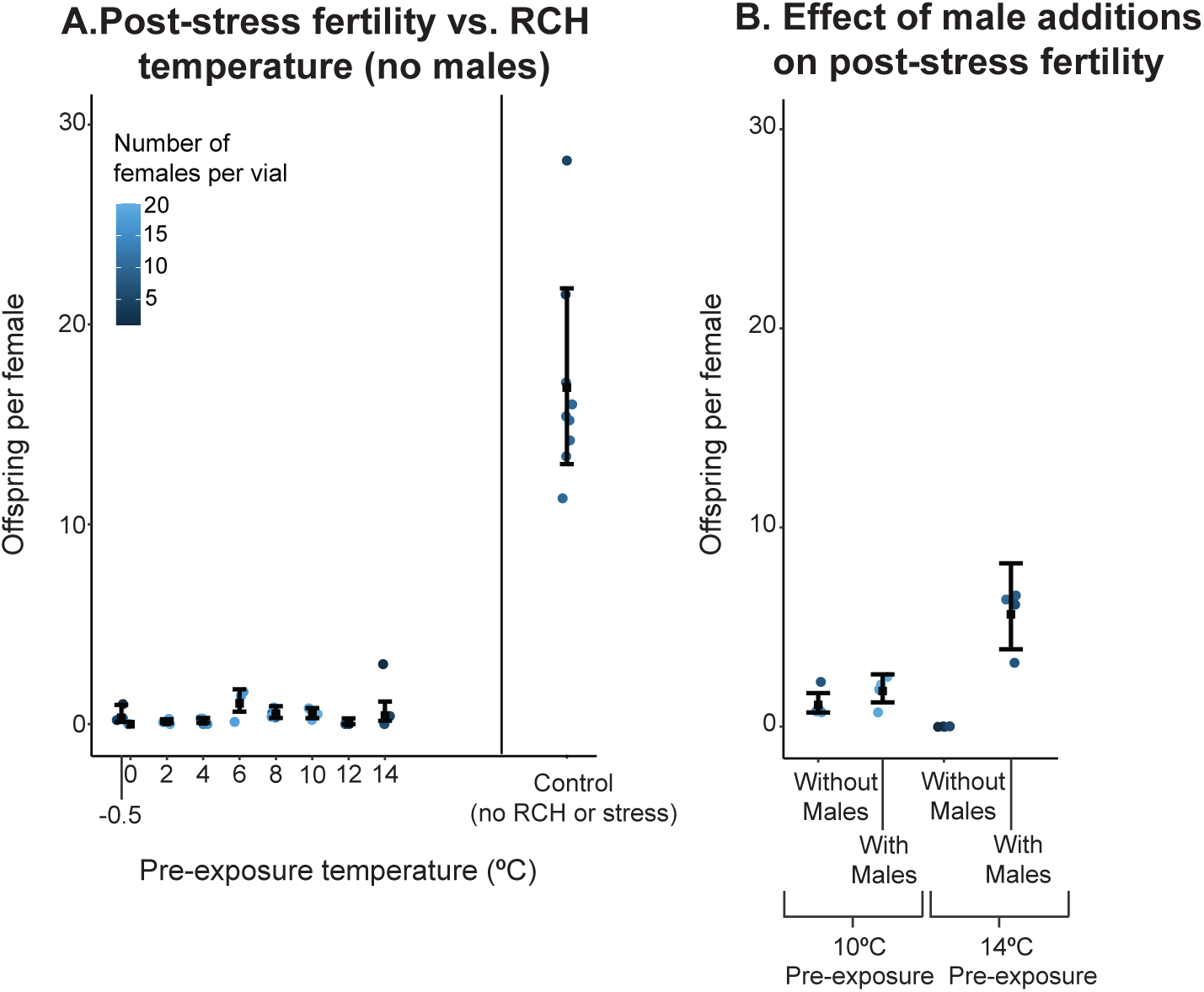
Effects of Pre-exposure temperature and male availability on female fertility. All flies were held at the pre-exposure temperature for 210 minutes followed by 60 minutes of stress exposure at -7°C as in the survival experiments. Each filled circle represents offspring per surviving female per replicate vial; post-stress, vials varied in the number of surviving females (blue shading). Black, filled squares represent the mean of each treatment estimated from fitted generalized linear models, while black whiskers represent 95% CI of the means. Panel A illustrates fertility of females without access to males post-stress compared to untreated controls. Panel B illustrates the combined effects of pre-exposure temperature and male addition on fertility. All replicate vials yielded zero offspring for 0°C pre-exposure (panel A) and 14°C pre-exposure without males (panel B) – these treatments thus do not have associated 95% CI.

Overall, pre-exposure to mildly cold temperatures allowed flies to successfully reproduce following cold shock in the lab. Estimated replacement rates (survival × offspring per female) following cold shock under laboratory conditions were low but clearly non-zero at most pre-treatment temperatures that allowed survival (Figure 5). Lab estimates of replacement rate were also highest at intermediate pre-exposure temperatures driven by higher survival resilience and slightly higher estimates of fertility resilience.

**Figure 5.**
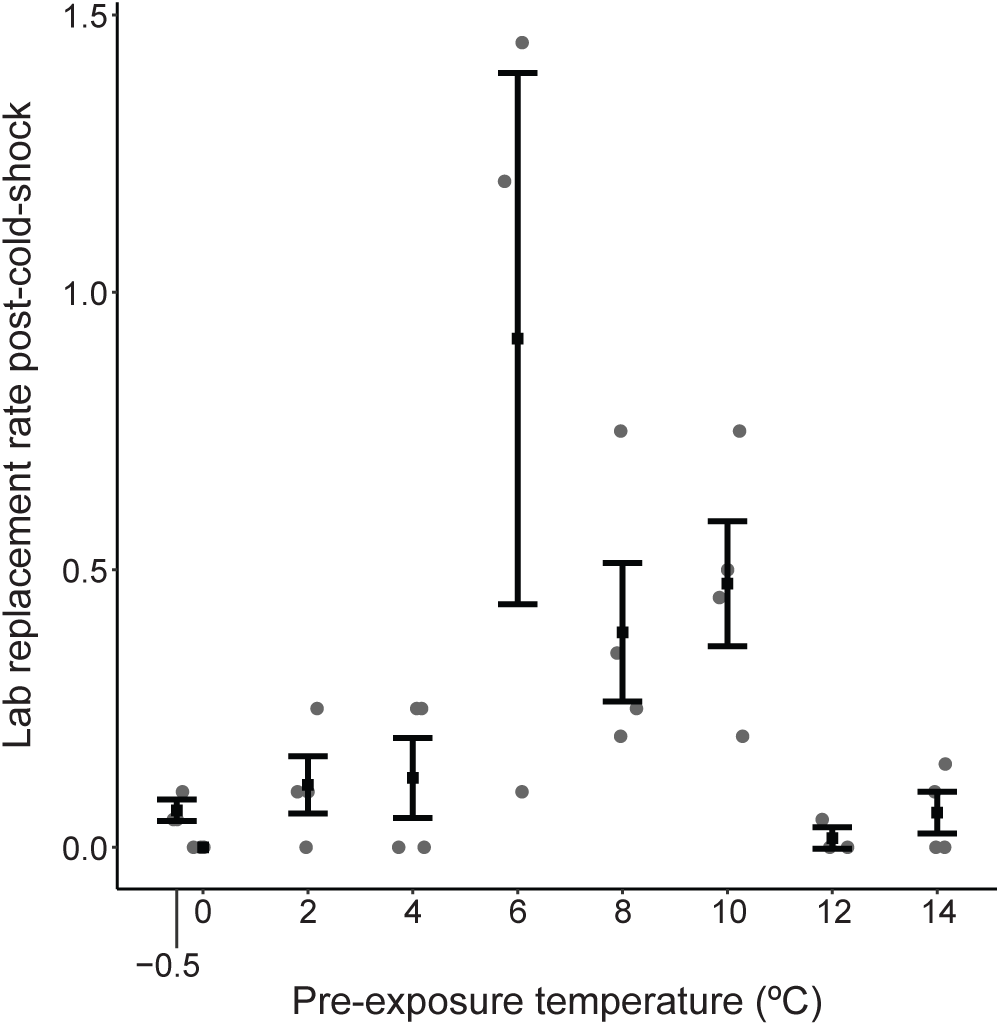
Replacement rate (survival proportion × offspring per female) estimated for each pre-exposure temperature in which both survival and fertility (pupa per female) were measured (excluding -1°C at which survival was zero; without addition of males post-cold shock). Each filled grey circle represents a replicate vial, filled black squares represent estimates of the mean, and error bars are approximate 95% CI (mean +-2SEM; except for 0°C pre-exposure at which no offspring were produced).

## Discussion

### Modeling cold pre-exposure as a hormetic process

The bi-logistic model that we implemented based on conventions for hormetic processes (Beckon et al., 2008) provides a useful quantitative framework to describe what appear to be sharp, threshold relationships between pre-exposure temperature and survival resilience. The model can also be fit to different distributions such as the negative binomial for fertility rates (Supporting file 1) though we used glms in our analysis of fertility. More flexible models could easily be applied to fit different curve shapes using other forms of the bi-logistic or other biphasic functions (Brain and Cousens, 1989; Hoekstra, 2006; Beckon et al., 2008), providing a flexible framework for understanding how acclimation and injury interact to influence phenotypes as pre-exposure temperatures approach lethal limits.

Here, our hormetic model fit is consistent with threshold processes for beneficial acclimation and cold injury separated by a relatively broad range of temperatures over which pre-exposure induces high survival resilience in *D. melanogaster*. The observation of a broad range of effective pre-exposure temperatures aligns well with the combined results of multiple studies suggesting that RCH (with respect to survival) occurs with sufficiently long pre-exposures (typically > 1h) at 0 - 5°C in *D. melanogaster* (Chen and Walker, 1994; Yi et al., 2007; Overgaard et al., 2007; Gerken et al., 2015). More remarkably, results from Chen et al. (1987) show a very similar plateau-shaped relationship in pharate adult *S. crassipalpis* (Fig. 3B). The cold hardening response seems to be at least partially cell autonomous (Nadeau and Teets, 2020), perhaps explaining the conservation of RCH across insect life stages and species (Teets et al., 2020). Concordance between the results of the current study and Chen et al. (1987) further suggests a broader conservation of the plateau-shaped, hormetic relationship implying temperature thresholds for hardening and cold injury.

### Limited protection of reproduction by cold pre-exposure

Fertility was low across all tested pre-exposure temperatures (about 85% lower than control flies) that enhanced survival resilience Nevertheless, some females successfully reproduced following cold shock when they were pre-exposed to more moderate cold. A previous study of cold hardening in *D. melanogaster* likewise showed fertility resilience to cold shock (-5°C for 1h) with cold pre-exposure (ramp down from 25°C to 0°C at -0.1°C/min followed by 1h at 0°C), yet fertility was substantially reduced compared to unstressed controls (Overgaard et al., 2007). In contrast, both the current study and Overgaard et al. (2007) found that RCH maintained survival at relatively high levels compared to unstressed controls, at least under some pre-exposure treatments. A study in *S. crassipalpis* also found more drastic reductions in fertility (about a 50% decline in fertilized eggs) compared to survival (Rinehart et al., 2000). There is one report that RCH can produce very high fertility resilience in *D. melanogaster*; pre-exposure to cold yielded post-cold shock fertility rates indistinguishable from unstressed controls (Kelty and Lee, 1999). However, that study applied a substantially less intense cold shock (-3°C for 1h, using the same Oregon R strain as the current study) and measured egg production but not viability through pupation. Overall, results from studies in *D. melanogaster* suggest that even when pre-exposures to cold lead to high survival resilience, extreme cold exposures may severely limit reproductive success regardless of temperature pre-conditioning.

### Ecological implications

Rapid cold hardening and similar experiments are often designed to simplify and isolate thermal effects, rather than to emulate ecological reality. However, rapid cold hardening can occur at ecologically relevant cooling rates that reflect, e.g., daily temperature fluctuations in the field (Kelty and Lee, 1999). Further, insects sampled directly from the field at different times of day also demonstrate cold tolerance consistent with RCH (Kelty, 2007).

Survival of fluctuations from moderate to extreme cold in nature surely reflect the offsetting effects of hardening and cold injury accumulating over time. Cold injury appears to accumulate at a constant rate over time at a given temperature, and that rate is negatively, log-linearly related to temperature in adult *Drosophila suzukii* (Tarapacki et al., 2021). This result is consistent with the thermal death time (TDT) models developed to predict heat tolerance in *D. melanogaster* based on a similar log-linear (positive) relationship between temperature and injury rate (Jørgensen et al., 2019). However, the TDT models predict survival under dynamic exposure to hot temperatures remarkably well under the assumption of no hardening (Jørgensen et al., 2019). In contrast, pre-exposure at moderately low temperatures has a strong, positive effect on survival resilience following a cold shock that initially accumulates over time, then eventually declines at longer exposure durations (Chen and Walker, 1994). Tarapacki et al. (2021) have also shown that long-term, egg-to-adult acclimation shifts the intercept of the log-linear relationship between temperature and cold injury rate.

Modeling the effects of dynamic temperatures on survival in cold temperatures will therefore require integrating the TDT model with models relating temperature to hardening effects. Our hormetic model shows that hardening effects are not log-linearly related to temperature – for example, the beneficial effect of pre-exposure on survival resilience is similar across the entire plateau of the hormesis curve. These results predict that sufficient time spent at any temperatures spanning the hormesis curve plateau should have similar hardening effects. Indeed, different cold ramping rates in RCH experiments can have equivalent effects on survival resilience in *D. melanogaster* even though flies in the different treatments experience different durations of exposure to different temperature ranges (Kelty and Lee, 1999). A model combining TDT and hardening effects would therefore need to include an additional component accounting for the non-linear relationship between temperature and hardening. Developing such a model would require reasonably large, factorial experiments applying a range of static temperatures and durations of exposure for predictions and dynamic temperature for validation similar to those described in (Jørgensen et al., 2019).

Fertility was also maintained across a broad range of pre-exposure temperatures, albeit at relatively low levels. In turn, our laboratory estimates of replacement rate were non-zero under most pre-exposure conditions, and highest within the same range of pre-exposure temperatures that maximized survival. Natural populations of flies may therefore persist in the face of severe cold stress in naturally fluctuating environments, provided that replacement rates are also non-zero in nature and eventually recover within or across generations (replacement rates below 1 lead to population decline). On the other hand, fertility may be the primary limiting factor for persistence in environments where bouts of extreme cold are relatively common. Though we did not test colder extreme temperatures (below -7°C), we suspect that there are ecologically plausible combinations of extreme cold and pre-exposure that maintain some fly survival but do not permit enough reproduction for population persistence in nature. A number of studies are consistent with this hypothesis, showing that thermal sensitivity of fertility better predicts geographic ranges and climate change vulnerability than the thermal sensitivity of survival (Walsh et al., 2019; van Heerwaarden and Sgrò, 2021; Parratt et al., 2021).

### Physiology – accumulation of RCH, then damage

What physiological processes might underlie the upper temperature threshold for RCH that we observed? There is sound evidence that several processes may contribute to RCH. These include adjustments to membrane fluidity, calcium ion concentration and calcium signaling, cryoprotectant accumulation, autophagy, and others that are well summarized in Teets et al. (2020). To our knowledge, there are no studies that systematically explore the relationship between these candidate mechanisms and temperature – rather, they are generally investigated at one or more time points during static temperature exposures. Studies applying a range of pre-exposure temperatures could provide robust additional evidence for a given candidate mechanism if survival resilience were correlated with some indicator metric (e.g., transcript or protein, ion, or metabolite abundance) across pre-exposure temperatures. Such experiments might also reveal whether different sets of physiological processes contribute to hardening at different pre-exposure temperatures.

We hypothesize that the decline in survival resilience at lower temperatures in the hormetic curve reflects the accumulation of cold injury. Cold temperatures will eventually inflict damage at low enough temperatures, but this is not sufficient evidence for cold injury at pre-exposure temperatures below 0°C in *D. melanogaster*. It is also possible that the physiological processes contributing to hardening simply become less effective or inhibited at lower temperatures. However, Chen and Walker (1994) show that the hardening effect (increased survival of a cold shock) of pre-exposure to 0°C and 4°C starts to exponentially decline at exposure durations that also cause declining survival without a cold shock. Though we cannot rule out the possibility that colder temperatures also inhibit hardening processes, cold injury very likely contributes to the logistic decrease in survival resilience at lower pre-exposure temperatures. As with hardening, experiments correlating candidate processes such as calcium ion homeostasis (MacMillan et al., 2015b) with survival resilience across temperatures could provide robust tests for a role in cold injury.

While somatic damage may be low across a wide range of pre-exposure temperatures, female reproductive organs and/or stored sperm are either damaged at pre-exposures below 14°C, or hardening processes are not effective at protecting reproductive tissues from cold shock. Exposure to temperatures from 4°C to 11°C can reduce fertility in *D. melanogaster*, but only at durations exceeding 24 hours (Mockett and Matsumoto, 2014). Thus, injury to stored sperm or female reproductive tissue seems unlikely at pre-exposures at and above 4°C in our experiment.

Rather, our results suggest that most pre-exposure temperatures that protect against somatic injury do not protect against injury to both stored sperm and female reproductive tissue. As little as 15 minutes at -5°C can kill all stored sperm in the female spermathecae and seminal receptacle of *D. melanogaster*, though cold shocked females can lay a few viable eggs that were previously fertilized (Novitski and Rush, 1949). It is therefore likely that our cold shock at -7°C for one hour killed all stored sperm, and that pre-exposure to moderate cold offered little, if any protection. Moreover, the few offspring that were successfully produced probably hatched from eggs fertilized prior to cold shock treatment. Novitski and Rush (1949) also note that, though stored sperm is killed during cold shock at -5°C, female fertility rebounds when they are mated to fertile males. We observed a minimal fertility rebound in females with access to males and with a pre-exposure to 10°C, and a higher, moderate rebound with a pre-exposure of 14°C, though both had substantially lower fertility compared to untreated controls. This suggests some damage to unfertilized eggs and/or reproductive tissues that persists at least 48h after cold shock. It is possible that damage is repaired and fertility rebounds to higher levels with more recovery time - other studies suggest fertility resilience when females are allowed to mate for five or more days following cold shock (Novitski and Rush, 1949; Kelty and Lee, 1999). Nevertheless, our results clearly demonstrate that pre-exposures that protect against mortality do not necessarily protect reproductive output.

### Summary

Our data show a hormetic curve with a broad range of pre-exposure temperatures that effectively improve cold shock survival, but yield universally low fertility. The fit of our biphasic model to these data suggests threshold processes underlying both RCH and cold injury, with little overlap in temperature range (cold injury does not seem to accumulate until past the threshold). Further investigation associating hormetic curves for traits like survival and candidate physiological processes may help elucidate the mechanisms underlying RCH and cold injury.

## Data availability

All code and raw data will be available on github and Zenodo upon publication in a peer reviewed journal.

## Acknowledgments

We would like to thank undergraduate researchers Zachary Courter and Gabrielle Dudek for their assistance with animal husbandry and Gage Tidwell for assisting with fertility experiments. We are especially grateful to Shelly McCain and James DeMayo for their feedback and support during the development of this research. This study was supported by NSF award number 2045263 to GJR.

